# The shape of the contact-density function matters when modelling parasite transmission in fluctuating populations

**DOI:** 10.1101/101535

**Authors:** Benny Borremans, Jonas Reijniers, Niel Hens, Herwig Leirs

**Affiliations:** Evolutionary Ecology Group, University of Antwerp, Belgium; Department of Ecology and Evolutionary Biology, University of California Los Angeles, United States; Interuniversity Institute for Biostatistics and statistical Bioinformatics (I-BIOSTAT), Hasselt University, Belgium; Department of Engineering Management, University of Antwerp, Belgium; Centre for Health Economics Research & Modelling Infectious Diseases (CHERMID – VAXINFECTIO), University of Antwerp, Belgium

**Author notes:** **Important notice** This is a pre-print version of the manuscript, made available through bioRxiv.org. Note that this manuscript has not yet been peer-reviewed, and has been submitted to a peer-reviewed journal.

**Keywords:** Contact rates, Disease invasion, Mass action, Nonlinearity, Threshold

## Abstract

Models of disease transmission in a population with changing densities must assume a relation between infectious contacts and density. Typically, a choice is made between a constant (frequency-dependence) and a linear (density-dependence) contact-density function, but it is becoming increasingly clear that intermediate, nonlinear functions intermediate are more realistic. It is currently not clear however what the exact consequences would be of different contact-density functions in fluctuating populations. By combining field data on rodent host (*Mastomys natalensis*) demography, experimentally-derived contact-density data, and laboratory and field data Morogoro virus infection dynamics, we explored the effects of different contact-density function shapes on transmission dynamics and invasion/persistence. While invasion and persistence were clearly affected by the shape of the function, the effects on outbreak characteristics such as infection prevalence and seroprevalence were less obvious. This means that it may be difficult to distinguish between the different shapes based on how well models fit to real data. The shape of the transmission-density function should therefore be chosen with care, and is ideally based on existing information such as a previously quantified contact- or transmission-density relationship or the underlying biology of the host species in relation to the infectious agent.

## Introduction

The transmission of infections can be sensitive to changes in population density, especially in the case of fluctuating wildlife populations [1–3]. When modelling disease transmission, the probability of encountering an infected individual is typically assumed to be either independent of (frequency-dependent) or linearly dependent on (density-dependent) population density [4]. Sexually transmitted infections are generally described using frequency-dependent transmission because the number of sexual contacts is assumed to remain constant, regardless of population density [5]. Infections that are transmitted through regular “every-day” contacts are often assumed to be density-dependent [6].

The choice of which contact-density function to use in a model of disease transmission entails potentially significant consequences. Most importantly, the two functions differ in whether or not an infection is expected to persist below a critical density of individuals [7]. The basic reproductive number (R_0_), defined as the number of secondary infections arising from the introduction of one infectious individual entering a completely susceptible population, is a central epidemiological measure that characterises the spread of an infection, and provides an immediate approximation as to how rapidly an infection can spread [8]. In its simplest form, *R_0_ = βN*, where *N* is population size and *β* is the transmission coefficient that consists of the rate *p* of becoming infected through contact with an infectious individual, multiplied by the contact-density function that equals *cN/A* (where *A* is area) when linear (density-dependent) and *c* when constant (frequency-dependent), and random homogenous mixing is assumed [9]. When transmission coefficient *β* changes with density, theory predicts a density below which the transmission rate is too low, causing *R_0_* to fall below 1 and the disease to disappear, whereas no persistence threshold density exists when *β* remains constant, independent of density [1,10,11]. Because changes in the transmission coefficient determine how quickly an infection can spread through a population, it can also be expected that the two contact-density functions will differently affect outbreak characteristics such as incidence, prevalence and outbreak size [12].

Because human populations are usually large and stable, many models of human disease transmission will not be significantly affected by the choice of the contact-density function. But when one is interested in modelling infection dynamics in populations of different sizes or periodically fluctuating densities, the shape of the assumed contact-density function may become highly important [3]. It is therefore not surprising that studies on how transmission rates and contacts relate to density have mainly been conducted in wildlife [9,13–17]. The main approach in these studies has been to measure disease prevalence in a field or experimental setting in which densities are manipulated or vary naturally, followed by fitting models with different transmission-density functions to the data.

There has traditionally existed a focus on whether transmission is frequency- or density-dependent, and rarely are other, nonlinear transmission-density functions investigated. For example, a study in which densities of the two-spot ladybird (*Adalia bipunctata*) were manipulated and transmission of sexually transmitted parasitic mite *Coccipolipus hippodamiae* was measured, only tested whether the transmission-density relationship was density-dependent or not, even though their results strongly suggest that the relationship more closely resembles a nonlinear asymptotic power function [18]. Similarly, a recent study in which strong emphasis was put on estimating a persistence threshold for Sin Nombre virus transmission in deer mice, assumed linear density-dependence without further investigation, although nonlinear transmission-density functions may have altered model results [19].

Nevertheless it has been well established that the binary distinction between density-independence or linearity can be inadequate, and a range of other possible nonlinear transmission functions has been suggested, often in the power law, asymptotic or logistic family [9,20–23]. Cowpox virus dynamics in a natural population of field voles (*Microtus agrestis*) for example has been shown to be best described by a nonlinear power function intermediate between density independence and linearity [13]. Intermediate density-dependence was also observed in *Ambystoma tigrinum* virus transmission in the tiger salamander [20].

Here, we want to investigate whether, and in which situations, implementation of the exact shape of the transmission response function is important. Although it would be possible to mathematically model the effects of different transmission-density functions for any hypothetical combination of demographic pattern and contact-density function, the sheer number of possible functions for each demographic situation would make it almost impossible to decide which functions are biologically relevant. To inform such models in a meaningful way we need biological background data, i.e. a species for which population dynamics, a contact-density or transmission-density function, and disease dynamics have been quantified, but until recently no such data were available. Using a combination of data from recent experiments in which we quantified contact rates across a wide range of population densities in the rodent *Mastomys natalensis* [24] and the infection parameters of Morogoro virus (MORV) in this rodent [25], we tested the effect of different transmission functions using a simple SIR transmission model in annually fluctuating host populations. By implementing a range of hypothetical combinations of infectious period, transmission rate and population size, we assessed what the effects of the contact-density would be for infections with different characteristics.

## Materials and methods

### Background data

Natal multimammate mice (*Mastomys natalensis*) occur throughout Sub-Saharan Africa, and are an important agricultural pest species and natural reservoir hosts for several microparasites that cause disease in humans, including *Yersinia pestis* (bubonic plague), leptospirosis and several arenaviruses including Lassa virus, which can cause severe haemorrhagic fever in humans [26–30]. Multimammate mouse demography has been studied thoroughly, which allows us to create a simple but accurate demographic model that will serve as a basis for transmission modelling (see below). In Tanzania, where most of the studies on its population ecology have been conducted, *M. natalensis* exhibits strong annual population fluctuations, with densities ranging from 10/ha in the breeding season to > 300/ha outside the breeding season [31,32]. Importantly, we have recently quantified a contact-density relationship for this species, which provides us with a realistic biological background for fitting the transmission functions [24].

The simulated infection dynamics in this study are based on those of Morogoro arenavirus (MORV), which naturally occurs in *M. natalensis* in Tanzania, and of which the transmission ecology [33] and patterns of infectivity [25,34] have been documented in detail; it has a latent period of about 3 days between infection and excretion (which we here ignored for simplicity), an infectious period of 30-40 days, and presumably lifelong immunity. MORV transmission can therefore be modelled using a simple SIR model (described below).

### Study design - models

We investigated the effect of four different contact-density functions using stochastic MORV transmission models. While transmission in these models is stochastic, demography was implemented deterministically because this allows us to focus entirely on the effects of stochasticity in transmission, and because it reduces computation time.

### Demographic model

The seasonally fluctuating densities of *M. natalensis* were modelled using a seasonal birth-pulse function, *B(t)* = *k* exp [–*s* cos^2^(*πt – φ*)], as described in Peel *et al*. [35]. This is a flexible function in which a synchrony parameter (*s*) determines the length of the birth period, and another parameter (*φ*) determines the timing of the birth period. Parameter *k* ensures that the annual population size remains the same, by compensating the number of births for the (constant) mortality rate *μ* [35]. Function parameters were fitted visually to a 20-year dataset of monthly population densities of *M. natalensis* in Tanzania ([31,32] and more recent unpublished data; Figure S1-1 in Supplementary Information). This deterministic demography was used as a basis for modelling demography in the stochastic SIR model (described below) in order to avoid the influence of stochastic changes in population density, and because this reduces the number of simulations that need to be performed.

### Transmission model

A standard SIR (Susceptible, Infectious, Recovered) model was used to simulate MORV transmission [36,37], described by the following set of coupled ordinary differential equations:

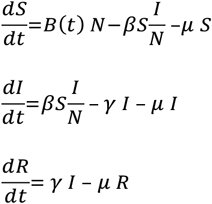

where *B(t)* is the time-dependent birth function described earlier, *μ* represents the mortality rate, *γ* is 1/infectious period, and *β = cp* is the transmission coefficient, which is composed of *p* (rate at which S becomes I when in contact with an I individual) and contact-density function 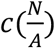 that can acquire different shapes depending on population density (explained below).

In order to compare invasion and persistence probabilities between different contact-density functions, we used a stochastic discrete time version of this SIR model, where the transition rates between categories were modelled stochastically, resulting in two possible stochastic events: infection (decrease of S, increase of I) and recovery (decrease of I, increase of R). Events were assumed to occur continuously in time at a certain rate, and were modelled using the “adaptive tau-leap” algorithm described in [35,38]. Briefly, each short time-step *δt*, the number of events of each type that occurs is randomly drawn from a Poisson distribution with mean *r_i_ δt*, where *r_i_* is the rate of each type of event *i*. If the number of simulated events would cause any of the categories (S, I or R) to fall below 0, *δt* is halved and new events are drawn (= “adaptive tau-leap”).

The model started at *t* = 0 and one infected individual was introduced after one year, at *t_0_ = 1 (I → 1)* in order to allow the initial population dynamics to stabilise. Different introduction times only had an effect on the linear and sigmoid functions, where they resulted in lower invasion probabilities between *t_0_* values of 1.2 and 1.4, which was likely a result of the low population densities (Figure S2-1 in Supplementary Information). There was no effect of introduction time on disease persistence (Figure S2-2 in Supplementary Information).

### Four different contact-density functions

The core of this study is the implementation of four different, biologically relevant contact-density functions 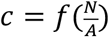 (Figure 1):

a. Constant function (or “frequency-dependence”) 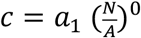, with fitting parameter *a_1_*. Independent of density, and typically (but not only) used in the case of sexually transmitted infections where the number of sexual contacts is not expected to change with population density [5,7].
b. Linear function (or “density-dependence”) 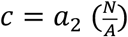, with fitting parameter *a_2_*. Typically used when assuming random mixing where (infectious) contacts increase linearly with population density [37,39].
c. Power function 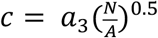, with fitting parameter *a_3_*. Has been used as an “intermediate” between frequency- and density-dependence [9,20]. Raised to a power of 0.5 in order to get the intermediate saturating shape. Contact rates increase at low densities, but the slope decreases at higher densities towards an asymptotic limit. This shape has been observed for contact rates in brushtail possums and elk [40–42], and has been shown to be a better predictor of cowpox transmission patterns than either frequency- or density-dependence [13].
d. Sigmoid function 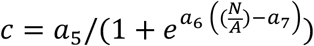 with fitting parameters *a_5_, a_6_* and *a_7_*. This function has a minimum, constant number of contacts at low densities, after which contact rates increase almost linearly with density until reaching a plateau when reaching a maximum number of contacts. This shape has been observed for multimammate mice contacts [24], and has been proposed previously as a biologically plausible function [9].

**Figure 1.**
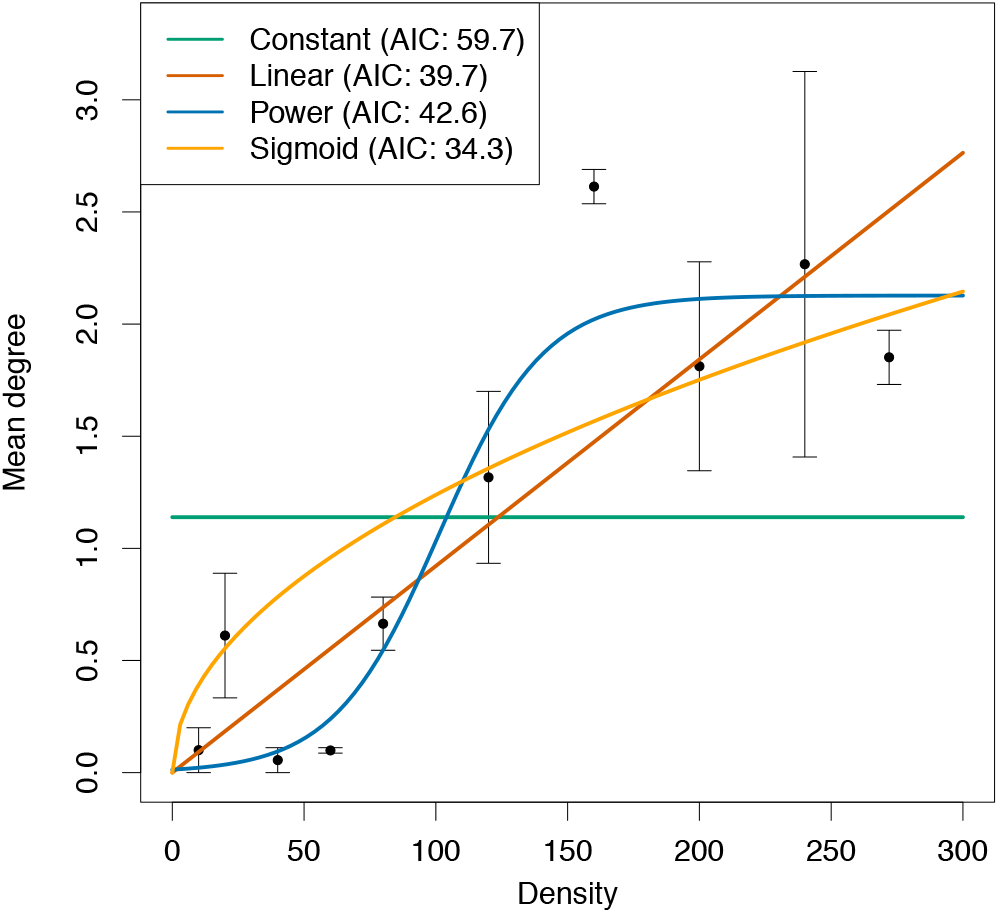
Contact-density functions fitted to experimental data from Borremans *et al*. [24], showing mean degree (the number of individuals one focus individual contacted) for a range of population densities (number of animals per ha = 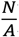).

These four different functions could in theory acquire an infinite number of shapes, so in order to realistically model them they were fitted to empirical contact-density data of *M. natalensis* [24,43] using maximum likelihood.

### Fitting transmission coefficient β

Considering that *β = cp*, after fitting contact parameter 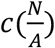 to contact-density data for the four functions, a transmission rate *p* had to be determined before being able to compare the effect of the different transmission-density functions. Equivalent to fitting model parameters to data, a function-specific constant (*q_i_*) had to be fitted for each function *i* to a common characteristic. Out of numerous possible characteristics to choose for fitting, we opted for one that ensured that *β*, summed across the probability distribution of population densities occurring during one year in a simulated, deterministic model of demography, was the same for each contact function. Formally, this meant that: 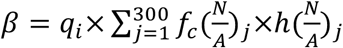, where *f_c_* is the contact-density function, *j* is population density and *h* is the frequency distribution of densities in a year. We chose this method because it has the advantage of not selecting for certain outbreak characteristics such as prevalence or outbreak size. We nevertheless also examined the effect of using two alternative fitting methods. The first alternative method fits *q_i_* so that a deterministic transmission model resulted in a maximum annual prevalence of 40%. The second alternative method fits *q_i_* so that the final annual number of infections was 2*N_0_*, i.e. twice (an arbitrarily chosen number) the initial number of individuals. While these alternative fitting methods resulted in highly different values for the constant (density independent) function, the three other functions were always very similar (Table S3-1 and Figure S3-1 in Supplementary Information). As we are mainly interested in the differences between the three non-constant functions, we did not further investigate the effects of these alternative *β*-fitting methods.

### Statistics

The effects of the contact-density functions were investigated through a number of meaningful epidemiological parameters: (1) SIR dynamics, including prevalence 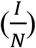; (2) Invasion probability, defined as the proportion of stochastic simulations in which the infection manages to survive the first year after introduction, conditional on having started successfully (successful start = infection persistence time > *t_0_* + infectious period); (3) Persistence probability, defined as the proportion of simulations in which, conditional on having survived the first year, the infection is still present at *t* = 10 years.

Invasion and persistence were estimated under a number of conditions of population size (*N_0_*), infectious period 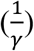 and transmission rate (*p*), where for each combination of these conditions 1,000 simulations were run. While we modelled changes in population *density* for each combination of parameters, we also assessed the effect of population *size* because this is expected to affect the probability for the infection to disappear from the population, independent of density. In order to ensure that we here implemented the effects of population size and not density, population density 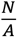 was calculated assuming that the area occupied when initial population size *N_0_* = 100 is 1 ha, and that area increases linearly with increasing values of *N_0_* (i.e. when initial population size increases, area also increases). Tested infectious periods: 1, 7, 14, 30, 60, 120 days. Tested transmission rates: 1, 5, 10, 25, 50, 75, 100, 125. Tested population sizes: 100, 500, 1000, 5000, 10000, 50000, 100000.

## Results

### Contact-density function fit to data

The four fitted contact-density functions were: 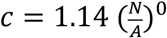 (constant), 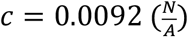 (linear), 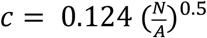 (power), 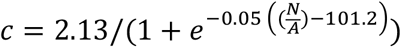 (sigmoid). As shown in Figure 1, a sigmoid contact-density function resulted in the best fit to contact-density data measured for *M. natalensis* in a previous field experiment [24], while the constant function clearly did not fit the data well. AIC values for the functions were 59.7 (constant), 39.7 (linear), 42.6 (power), 34.3 (sigmoid). Akaike weights were 0.00 (constant), 0.06 (linear), 0.01 (power), 0.92 (sigmoid), providing additional support for the much better fit of the sigmoid function.

### SIR dynamics and prevalence

Because we are interested in the qualitative effects of the contact-density functions rather than in detailed differences that are specific to the model system, we here report the results with a focus on the qualitative aspects of SIR dynamics and prevalence. The constant function resulted in a relatively low epidemic peak during the breeding season, while the other three functions showed a clear epidemic peak where prevalence (proportion Infecteds; Figure 2, red curves) increased sharply for the linear and sigmoid functions, but more gradually for the power function. Median peak prevalence for the four functions: 45% (constant), 68% (linear), 60% (power), 72% (sigmoid). The dynamics of the proportion of Recovered (immune – antibody-positive; Figure 2, blue curves) individuals was highly similar for the four functions, although they did differ in how low the Recovered proportion becomes at the end of the birth pulse: 15% (constant), 2% (linear), 14% (power), 0.5% (sigmoid). There was a significant build-up of Susceptibles (Figure 2, green curves) for the linear and sigmoid than for the other two functions, due to the fact that transmission rates increased later in the year (at higher densities).

**Figure 2.**
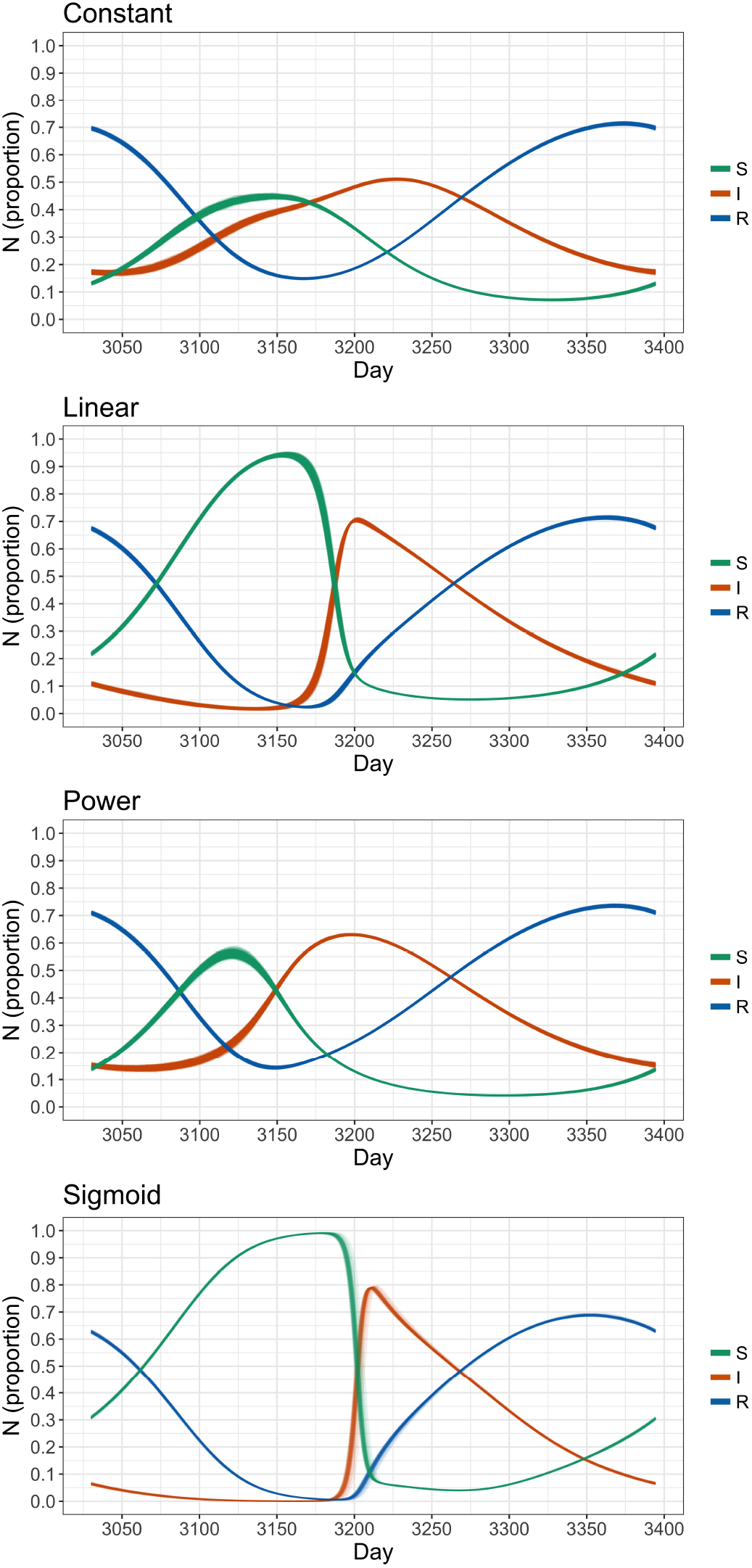
SIR dynamics for the four different contact-density functions. For each simulation run in which there was successful persistence, the 8^th^ year was retained (days 3030 to 3395). The figure shows all these outputs plotted on top of each other. The increase in susceptibles corresponds with the birth pulse. Infectious period 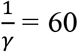 days, transmission rate *p* = 50, initial population size *N_0_* = 100,000.

### Invasion and persistence

Invasion and persistence probabilities were investigated for a range of population sizes (*N_0_*), infectious periods (1/*γ*) and transmission probabilities (*p*). Note that while infectious period results are reported in absolute days, they can also be interpreted in relation to the demographic timescale used in the simulations (e.g. annual breeding, brief recruitment period), which will aid comparison with other pathogen-host systems in which host densities fluctuate [35].

Successful invasion and persistence were more often observed for the constant function than for the other functions (Figures 3 and 4). Even at low population sizes successful invasion was almost certain for infectious periods of 30 days and longer, and was even observed for an infectious period of 7 days in sufficiently large populations. In contrast, for the other functions successful invasion was never observed below infectious periods of 7 days, and even with an infectious period of 30 days invasion was rare for the sigmoid function. For the linear function, invasion probabilities were lower than those for the power function, whereas persistence probabilities for these two functions were similar. The sigmoid function resulted in the lowest invasion or persistence success, where persistence was rarely observed for an infectious period of 30 days. Even when the infectious period was 60 days, persistence was only observed when population size was sufficiently large (e.g. 50% for a population size of 5000).

**Figure 3.**
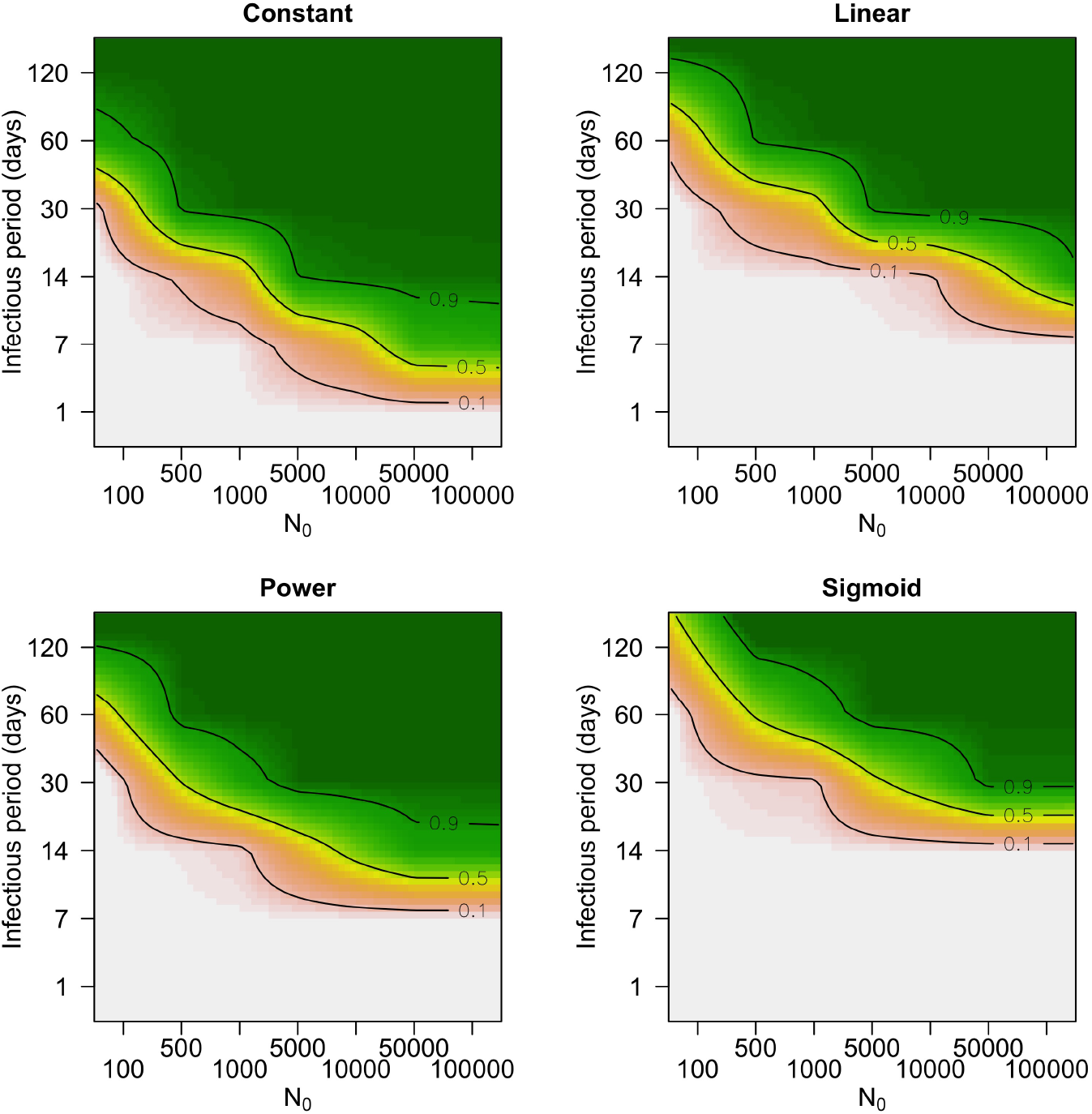
Invasion probabilities for the different contact-density functions, for a range of infectious periods 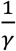 and initial population sizes *N_0_* (transmission rate *p* = 50). Simulations were conducted for all values indicated by tick marks on the axes, and results are interpolated between these values for illustration.

**Figure 4.**
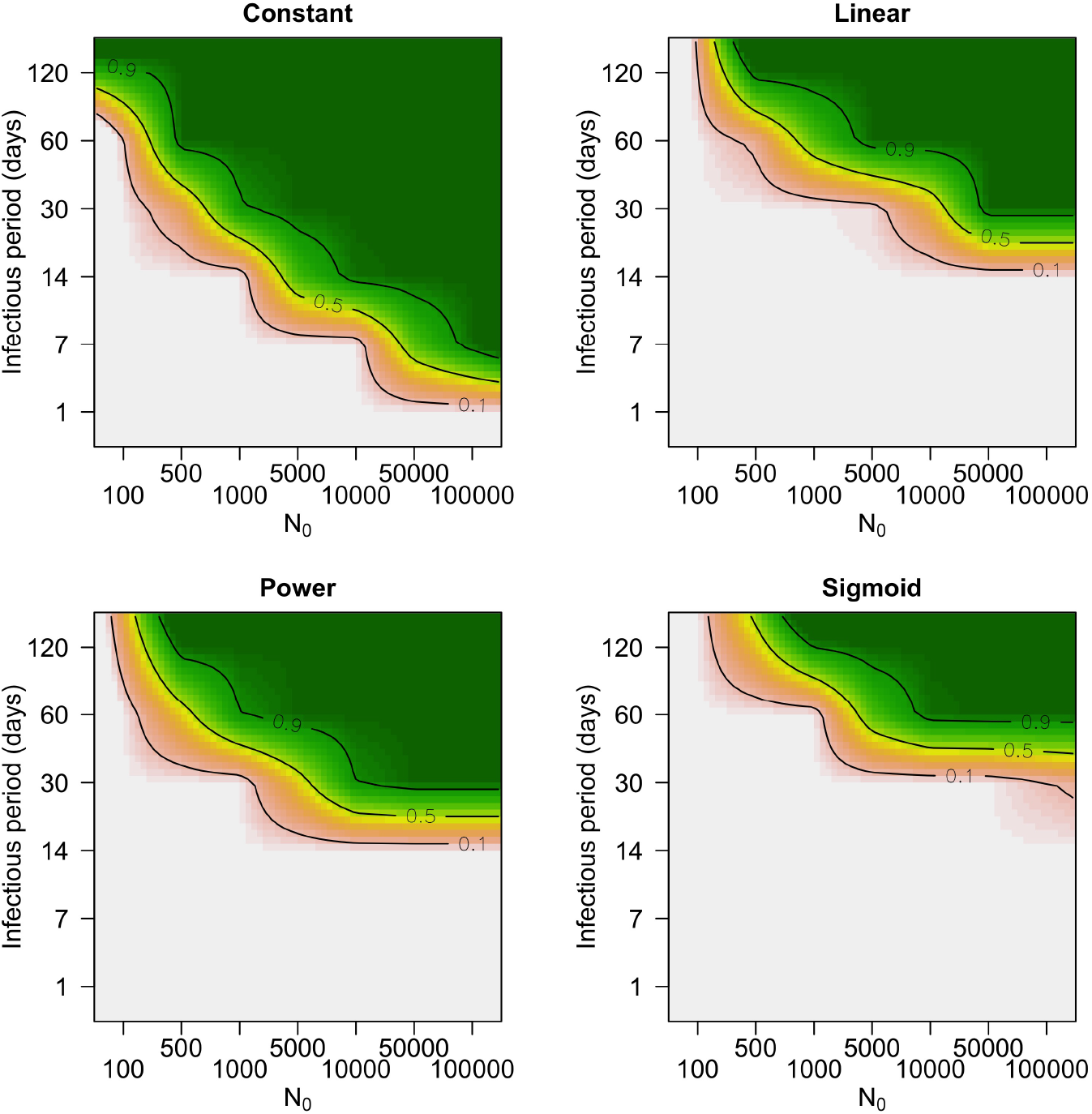
Persistence probabilities for the different contact-density functions, for a range of infectious periods 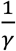 and initial population sizes *N_0_* (transmission rate *p* = 50). Persistence probabilities were calculated using only simulation runs in which there was successful invasion. Simulations were conducted for all values indicated by tick marks on the axes, and results are interpolated between these values for illustration.

Invasion, but not persistence, was affected by the time of the year at which the infection was introduced (Figures S2-1 and S2-2 in Supplementary Information). For *t_0_* = 1.2 and *t_0_* = 1.4, which corresponds with the low-density period (Figure S1-1 in Supplementary Information), invasion probability was generally lower than for other introduction times.

The effect of transmission rate on invasion and persistence were similar between the constant, linear and power functions, although the constant function still resulted in more successful invasion/persistence at lower transmission probabilities (Figures S4-1 and S4-2 in Supplementary Information). The sigmoid function however was more affected by transmission probabilities than the other three functions, where for an infectious period of 30 days persistence was only possible for a combination of high transmission rate and large population size (Figure S4-2 in Supplementary Information).

## Discussion

Models of disease transmission in which population densities change cannot avoid assuming a way in which transmission scales with density [9]. Using data-driven models of virus transmission in a population with seasonal births, we found that the shape of the transmission-density function can have important consequences for invasion and persistence success.

While SIR dynamic patterns for the constant function were distinct from those of the three density dependent functions, these three functions did not result in highly different SIR dynamics. Infection prevalence (the proportion if Infecteds) and seroprevalence (the proportion of Recovereds) patterns were very similar for the linear and sigmoid functions. For the power function, seroprevalence and infection prevalence patterns were slightly smoother, with less pronounced peaks, than for the linear and sigmoid functions. Invasion and persistence probabilities were clearly affected by the shape of the transmission-density function. They were always higher for the constant function than for the other functions. The sigmoid function resulted in the lowest invasion and persistence probabilities, and was the most sensitive to population size, length of the infectious period and transmission rate. Invasion and persistence success for the linear and power functions were intermediate between the constant and sigmoid functions. Depending on the time at which the infection is introduced, the differences in invasion and persistence probability between the contact functions can become even more pronounced.

The different consequences of the contact-density functions can likely be attributed to a number of key differences in their shapes. Considering that the infection is most sensitive to extinction during periods of low population density, the size of transmission coefficient *β* at low densities will be a highly influential factor. An important consequence of this is that larger population sizes are necessary for successful disease invasion/persistence when *β* is low during low-density periods. In our case, for example, a minimum population size of 10,000 (equivalent to a 100ha area) was necessary for a 50% persistence success rate for the power function (30-day infectious period), while this was 50,000 for the linear function and larger than 100,000 (not tested) for the sigmoid function. Knowledge of contact rates at low population densities is therefore critical when estimating invasion and persistence thresholds. A second important factor is the rate at which *β* increases with density. The epidemic peak will be more pronounced when there is a strong increase of *β* with density (e.g. the sigmoid function at intermediate densities, but also the linear and power functions). The sigmoid function for example results in a steep increase in transmission rates during the juvenile recruitment season as soon as a threshold density of susceptibles is reached (here around 80-100/ha).

When considering both the SIR dynamics and invasion/persistence of the three density dependent functions, a contrast emerges than can have significant implications for fitting a *β*-density function to epidemiological data: while there was a clear effect of function shape on invasion and persistence, SIR dynamic patterns were less distinct. This means that it could be difficult to discern between different contact-density functions when fitting model parameters to real, inherently noisy data [3,9,13]. Nevertheless, because the functions do introduce different invasion and persistence probabilities, it will in some situations be crucial to implement the correct function. Ideally this choice is based on the quantified contact-density or transmission-density relationship of the host/infection system that is being studied, but such data are rarely available.

A possible solution for this problem could be to establish/utilise general links between certain biological traits and contact and transmission patterns [44]. The shape of the transmission-density function is determined by a combination of infection and host characteristics, so based on these characteristics, it should theoretically be possible to *a priori* predict the shape of the function, at least roughly. Knowledge of density-dependent changes in home range size and overlap could for example be a useful proxy for the contact-density function. For male brushtail possums (*Trichosurus vulpecula*) it has been established that contacts increase with density according to a positive power function [41], which fits with the fact that this species is not territorial, and with the observation that home ranges are larger at low densities which may result in the maintenance of contacts. Such an inverse correlation between home range size and density was also observed for *M. natalensis* [45], and this may have similar results on contact rates at low densities, as maintenance of contacts even at very low densities was also observed for this species [24]. This pattern would be expected to be different for territorial species. In an enclosure experiment, movements of meadow voles (*Microtus pennsylvanicus*), which are strongly territorial, decrease significantly with density [46], and although the effect of density on contacts was not measured, it is not unlikely that this decrease in movement distance corresponds with a contact-density function that does not increase, or at least not linearly. As a final example, consider the experimental study of the transmission of the parasitic mite *Coccipolipus hippodamiae* in populations of the two-spot ladybird (*Adalia bipunctata*) [18]. The mite is transmitted sexually, and although sexual contacts are typically assumed to be frequency-dependent, the authors observed that the transmission-density function was closer to linear density-dependence and therefore concluded that the common assumption that sexual transmission is frequency-dependent is not always true. Their study species (*A. bipunctata*) however is known to be highly promiscuous, which means that sexual contacts are not limited to one or a few mates, but instead increase with density. *A priori* use of this knowledge about host and infection biology would have resulted in the more accurate prediction that sexual transmission of *C. hippodamiae* is density-rather than frequency-dependent.

Many wildlife species experience seasonal birth pulses and density fluctuations, and while it has been established that birth pulses can have strong effects on disease transmission [35], we now see that the shape of the transmission-density function can have further significant effects on disease invasion and persistence. The implementation of the transmission-density function should therefore be done with care, and as informed as possible. Although currently few studies have quantified the relationship between contacts and density, other relevant knowledge about host biology and behaviour can potentially be used for deciding on the best possible shape of the transmission-density function.

## Ethics statement

No experiments were carried out during this study.

## Data accessibility

Field data used in this study were reported previously in [24], and are freely available at [43]. Code used in this article for model simulation can be found as Supplementary Information.

## Authors’ contributions

BB, JR and HL conceived of the study; BB performed the analyses and wrote the first draft; All authors coordinated the study and helped draft the manuscript; All authors gave final approval for publication.

